# Glendonite occurrences in the Tremadocian of Baltica: first Early Palaeozoic evidence of massive ikaite precipitation at temperate latitude

**DOI:** 10.1101/486779

**Authors:** Leonid E. Popov, J. Javier Álvaro, Lars E. Holmer, Heikki Bauert, Mansoureh Ghobadi Pour, Andrei V. Dronov, Oliver Lehnert, Olle Hints, Peep Männik, Zhifei Zhang, Zhiliang Zhang

## Abstract

The Tremadocian (Early Ordovician) is currently considered a time span of greenhouse conditions with tropical water surface temperature estimates, interpolated from oxygen isotopes, approaching 40°C. In the high-latitude Baltoscandian Basin, these data are in contrast with the discovery of glendonite, a pseudomorph of ikaite (CaCO_3_·6H_2_O) and valuable indicator of near-freezing bottom-water conditions. The massive precipitation of this climatically sensitive mineral is associated with transgressive conditions and high organic productivity. Surprisingly, the precipitation of glendonite is contemporaneous with the record of conodonts displaying low δ^18^O values, which would suggest high temperatures (>40°C) in the water column. Therefore, the early Tremadocian sediments of Baltoscandia contain both “greenhouse” pelagic signals and near-freezing substrate indicators. This apparent paradox suggests both the influence of isotopically depleted freshwater yielded by fluvial systems, and the onset of sharp thermal stratification patterns in a semi-closed basin, which should have played an important role in moderating subpolar climates and reducing latitudinal gradients.

---

Except one rather controversial note^1^, the record of glendonites displays an apparent gap from Neopoterozoic^2^ to Permian^3^ times. However, similar calcareous nodular aggregates embedded in Tremadocian black shales of the East Baltica, so-called “antraconites”, have been known for more than 150 years. These aggregates are documented from 24 geographical localities in the Türisalu and Koporiye formations (*Cordylodus angulatus* - *Paltodus deltifer pristinus* zones) exposed along 600 km of the Baltic-Ladoga Glint^4^ and sporadically in the Orasoja Member (Kallavere Formation). All these units were accumulated in the Baltoscandian Basin (Fig. 1a), an epeiric sea with a central flat-floored depocentre rimmed to the south (recent coordinates) by a chain of low islands and associated shoal complexes^5,6^. During Tremadocian times, the basinal depocentre^4,7^ recorded black shale deposition episodically punctuated by wave and storm-induced processes, pointing to a sediment-starved, distal offshore-dominant clayey substrate, in which organic matter and trace metals became highly concentrated due to extremely low deposition rates and an exceptionally low influx of siliciclastic material^8^. In contrast, nearshore environments comprised uncemented, well-washed, cross-laminated quartzose sands, which included high concentrations of allochthonous obolid coquinas that were continuously reworked along the shorelines. The major Furongian-Tremadocian sea-level fluctuations are reflected in distinct stratigraphic gaps (Supplementary Figs 2, 4), some of them highlighted by the evidence of palaeokarst^24^.

**Figure 1.**
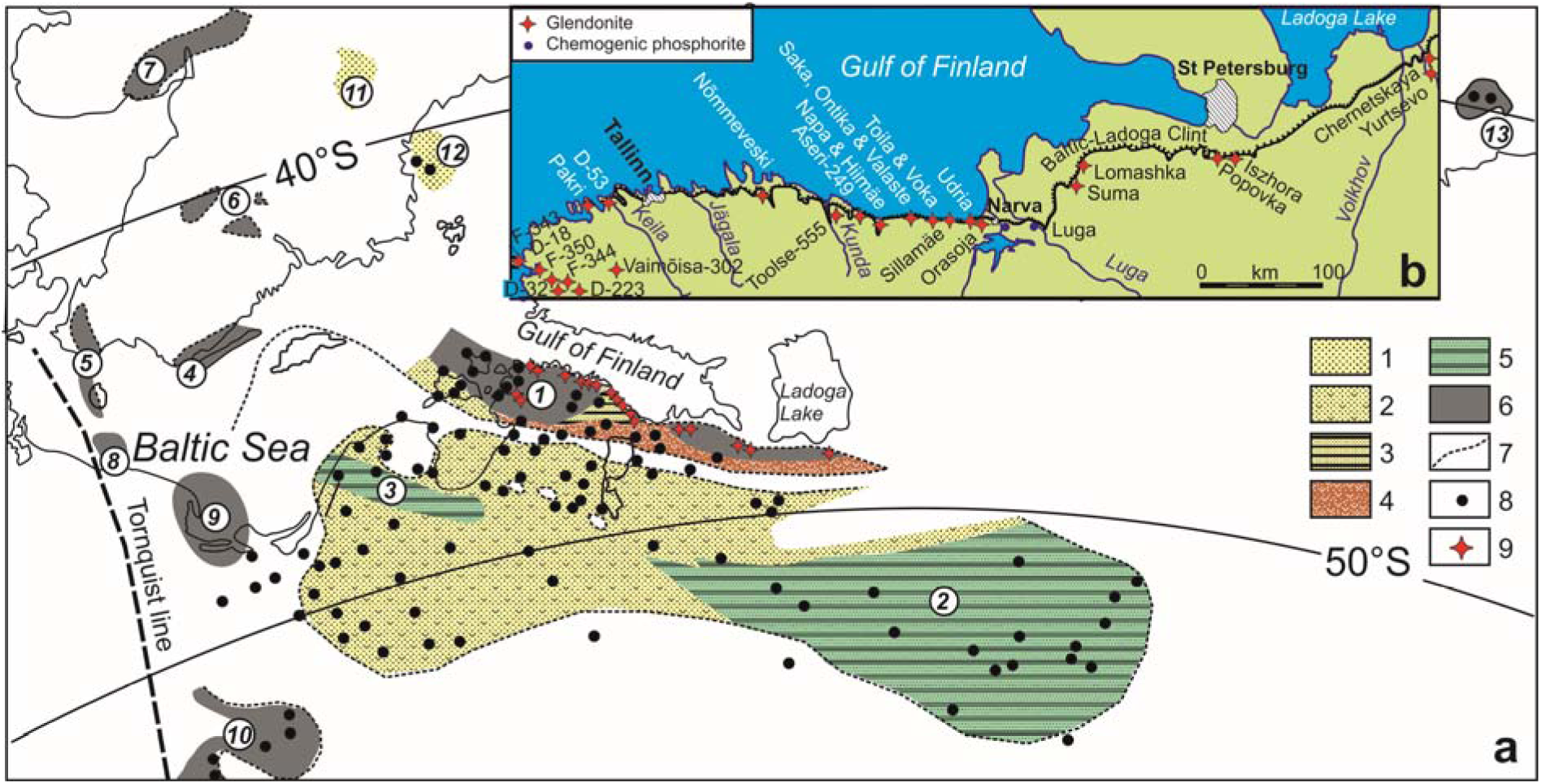
Distribution of major Tremadocian facies in the northern and central part of East Baltica with setting of glendonite localities; **(a)** facies map (after^5–6^) showing: 1, quartzose sands and sandstones (shoreface); 2, quartzose sands with obolid brachiopods (shoreface); 3, fine-grained sand and black shale interbeds (offshore); 4, detrital sands with abundant obolid brachiopods (beach and bar systems); 5, fine-grained sandstones, siltstones and claystones (offshore to basin); 6, black graptolitic shales (basin); 7, area of recent development of Tremadocian strata; 8, bore-holes; 9, glendonite localities; geographical areas: *1*, North Estonia and St Petersburg region; *2,* Moscow basin; *3,* Jelgava depression; *4,* Öland and Småland; *5*, Scania-Bornholm; *6,* Östergötland-Närke-Västergötland; *7,* Oslo Region; *8,* Łeba area; *9,* Łeba-Gdaǹsk area; *10,* Podlasie depression; *11*, Siljan, *12,* Bothnian Sea; *13,* Kolguev Island; palaeolatitudes for Tremadocian after^32^; (**b**) schematic map of the Baltic-Ladoga Clint area showing position of glendonite and phosphorite localities.

## Updated significance of Tremadocian antraconites

Antraconitic aggregates were sampled in the Tremadocian Türisalu Formation of North Estonia and the Koporiye Formation of the eastern St Petersburg region (Fig. 1 and Supplement Figs 2–4). Antraconites occur as single crystal pseudomorphs (Fig. 2g), stellate (Fig. 2e, h) and rosette clusters (Fig. 2b, c), usually 5–10 cm across and up to 20 cm in length encased in black shales. One characteristic glendonite horizon, 15 cm thick, embedded in a distinct grey graptolitic clay of the Koporiye Formation, is traceable over 3.5 km on the eastern bank of the Syas River, between Chernetshkaya and Yurtzevo villages^5^ (Fig. 2b). It represents a compact aggregate of crystals, with randomly oriented long axes on the top of a pyritised sandstone layer (Fig. 2f). The precipitation of ikaite probably occurred at the sediment-water interface, highlighting the top of a condensed bed.

**Figure 2.**
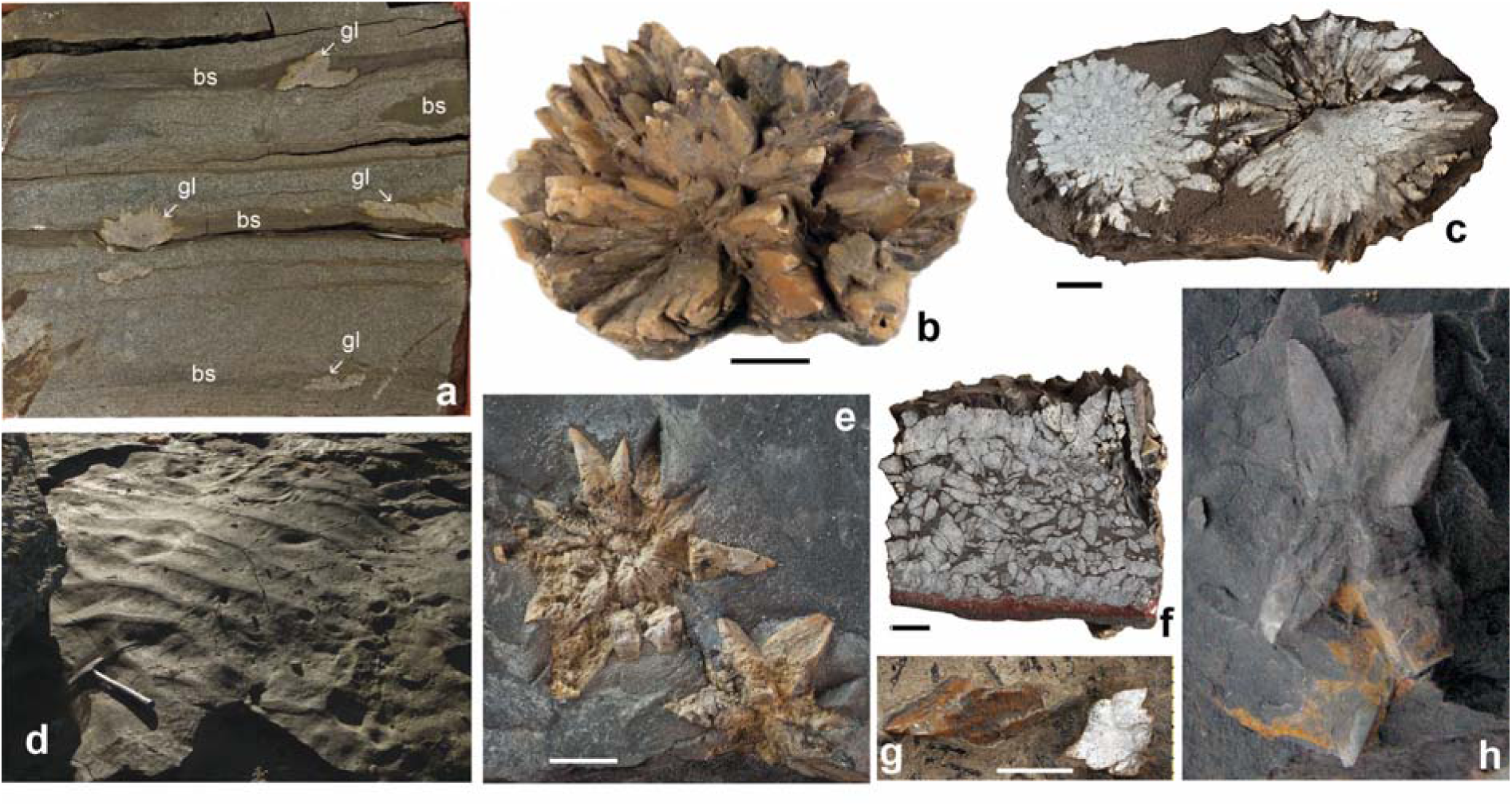
Photographs of glendonites and sedimentary features from the East Baltic black shales; (**a**) GIT 366-296, cut of a drill core showing silts and silty sandstones with thin layers of black shale (bs) bearing glendonite pseudomorphs (gl), D-223 borehole; (**b**) rosette cluster of glendonite pseudomorphs, Toolse Member (*Paltodus deltifer pristinus* Zone), Napa, Estonia; (**c, f, h**) cross sectional view of two rosette glendonite pseudomorph clusters, cross sectional view of “glendonite bed”, and stellate glendonite pseudomorph cluster, Koporiye Formation (*Cordylodus angulatus* Zone), Syas River near Yurtzevo, Russia; **(d)** bedding surface of black shale with wave generated symmetrical ripple marks, Tabasalu Member (*Cordylodus angulatus* Zone), Pakri Cape; **(e)** GIT 571-19, two stellate glendonite pseudomorph clusters, Tabasalu Member (*Cordylodus angulatus* Zone), D-32 borehore, Vidruka; **(g)** TUG 220-77b, blade shaped glendonite pseudomorph, Tabasalu Member, F-350 borehole, Kirimäe; (**b, d-f**) scale bars are 1 cm; (**e, g**) scale bars are 5 mm; (**a-g**) photo by H. Bauert; (**h**) photo by A. Dronov.

Glendonite crystals display a slightly distorted, prismatic habit, grading both laterally and centripetally into mosaics of microgranular calcite (Fig. 3a, b), commonly red-stained by dispersed ferroan oxi-hydroxides and variable content in organic matter. XRD analysis reveals a predominance of calcite, locally contaminated by silty quartz and feldspar and the scattered presence of dolomite rhombs. Cathodoluminescence (CL) petrography distinguishes rhombohedral, fine-fibrous, chevron-like, ovoidal, spherulitic and clear mosaics of calcite (Fig. 3c). Crosscutting relationships between different calcite phases suggest multiple stages of dissolution-recrystallization and (either complete or partial) replacement of pre-existing types of calcite. Successive recrystallization phases from an ikaite precursor is marked by non-luminescent to dull brown-orange (spherulitic) and reddish (fine-fibrous and mosaics) for younger neomorphic calcite mosaics. CL patterns reflect a progressive increase in luminescence of the youngest calcite generations.

**Figure 3.**
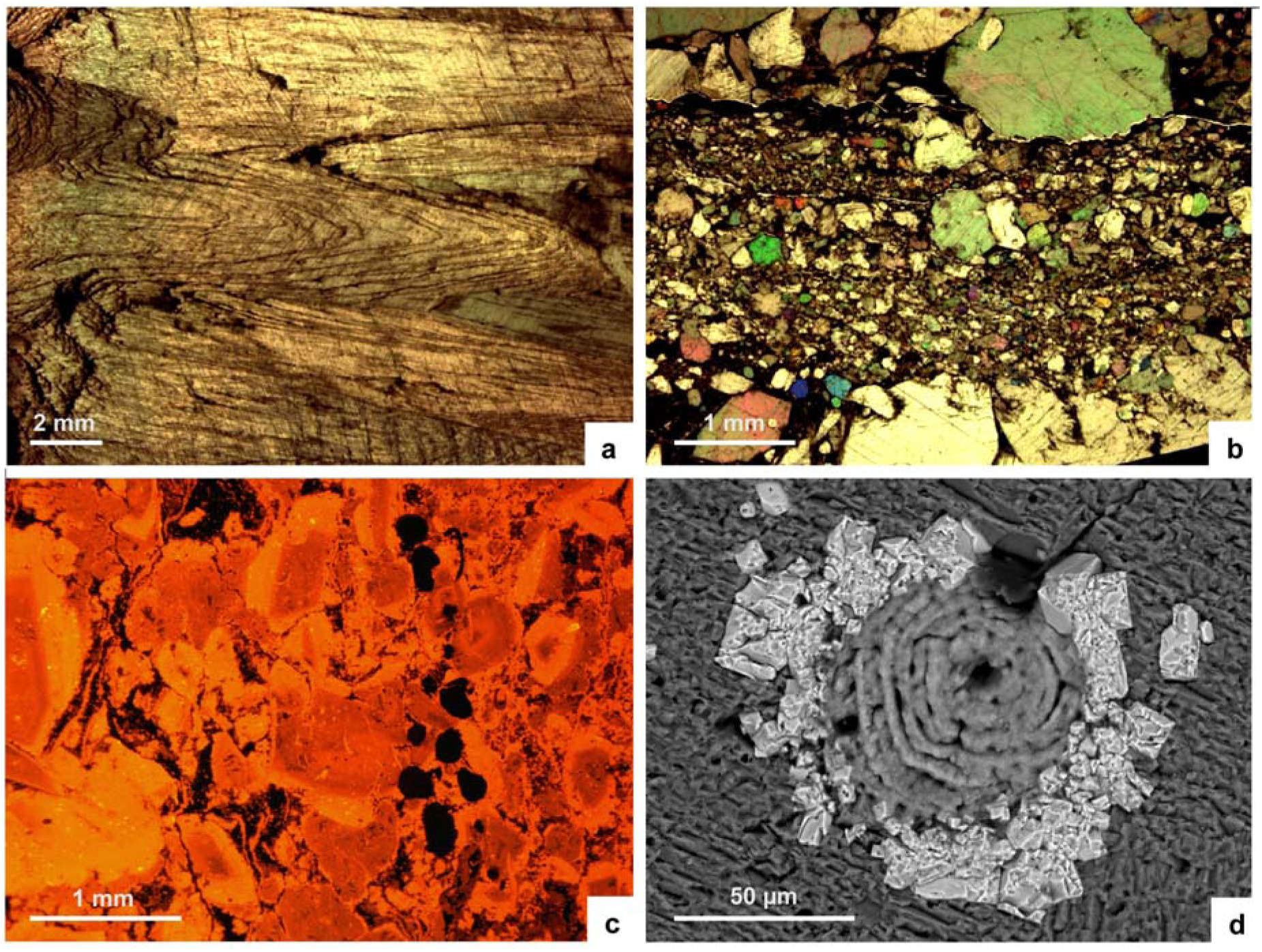
Thin-section photomicrographs of glendonite and associated sediment; **(a)** core of glendonite displaying the prismatic chevron-like habit of stellate pseudomorphs, Koporye Formation of St Petersburg region (petrographic microscope, plane light); **(b)** glendonite showing mosaics of microgranular calcite after recrystallization of original ikaite (petrographic microscope, parallel light), Türisalu Formation, North Estonia; **(c)** cathodoluminescence image of previous picture showing ovoidal calcite grains with red-CL colours surrounded by subsequent orange-CL overgrowths; **(d)** SEM image showing euhedral crystals of pyrite nucleated on external wall of a phosphatic linguliform brachiopod bioclast, Koporiye Formation of St Petersburg region.

The wrapping of shale laminae around the glendonites, which are surrounded by unconsolidated clays, demonstrates that the concretions and nodules lithified during early diagenesis. However, competency differences caused by variable early-lithification of crystal aggregates and interbedded claystone contributed to fracturing. Two kinds of fissures can be distinguished. Those unrecognizable under CL are up to 2 mm wide, have no preferential alignment and are commonly branched and anastomosing; walls are highly irregular and show a very poor (or absent) fit. These fractures formed in a soft, semi-cohesive substrate (Fig. 2a, c). In contrast, fissures easily detectable under CL, up to 1cm wide, have their walls dominantly composed of straight segments that exhibit a clear fit. Their porosity was occluded by a CL bright yellowish cement. They formed at a late stage of diagenesis in fully lithified concretions.

The dominant form of dolomitization occurs as euhedral to subeuhedral, dolomite rhombs, up to 60 μm in size. Locally, pervasive dolomitization has led to equigranular mosaics with little trace of primary textures. Fine-grained pyrite, identifiable as cubic, anhedral and framboidal forms and up to 8 μm in size, are locally abundant throughout the concretions and the encasing matrix. Occasionally, pyrite directly encrusted previous hard substrates, such as scattered fossil skeletons (Fig. 3d).

## Isotope data from glendonites

Carbon and oxygen isotope analyses were carried out in the glendonitic calcite mosaics in order to characterize the earliest diagenetic phases. The results are presented in Repository Data and graphically shown on Fig. 4. Isotopic data cluster in a single field, where carbon isotope ratios range from +0.6 to −8.9‰ and oxygen values from −5.7 to −8.2%o, and are consistent with isotope data derived from other methane-free glendonites^3, 9–11^. Modern glendonites typically show a much broader range in carbon isotope values (from +10 to − 40‰ ^3 10–13^ (Fig. 4), which are strongly dependent on the depositional environment: the extremely negative δ^13^C values (< −20‰) of many deep marine glendonites are likely controlled by the input of biogenic methane, whereas ikaite precipitated in lacustrine environments exhibits positive δ^13^C values.

**Figure 4.**
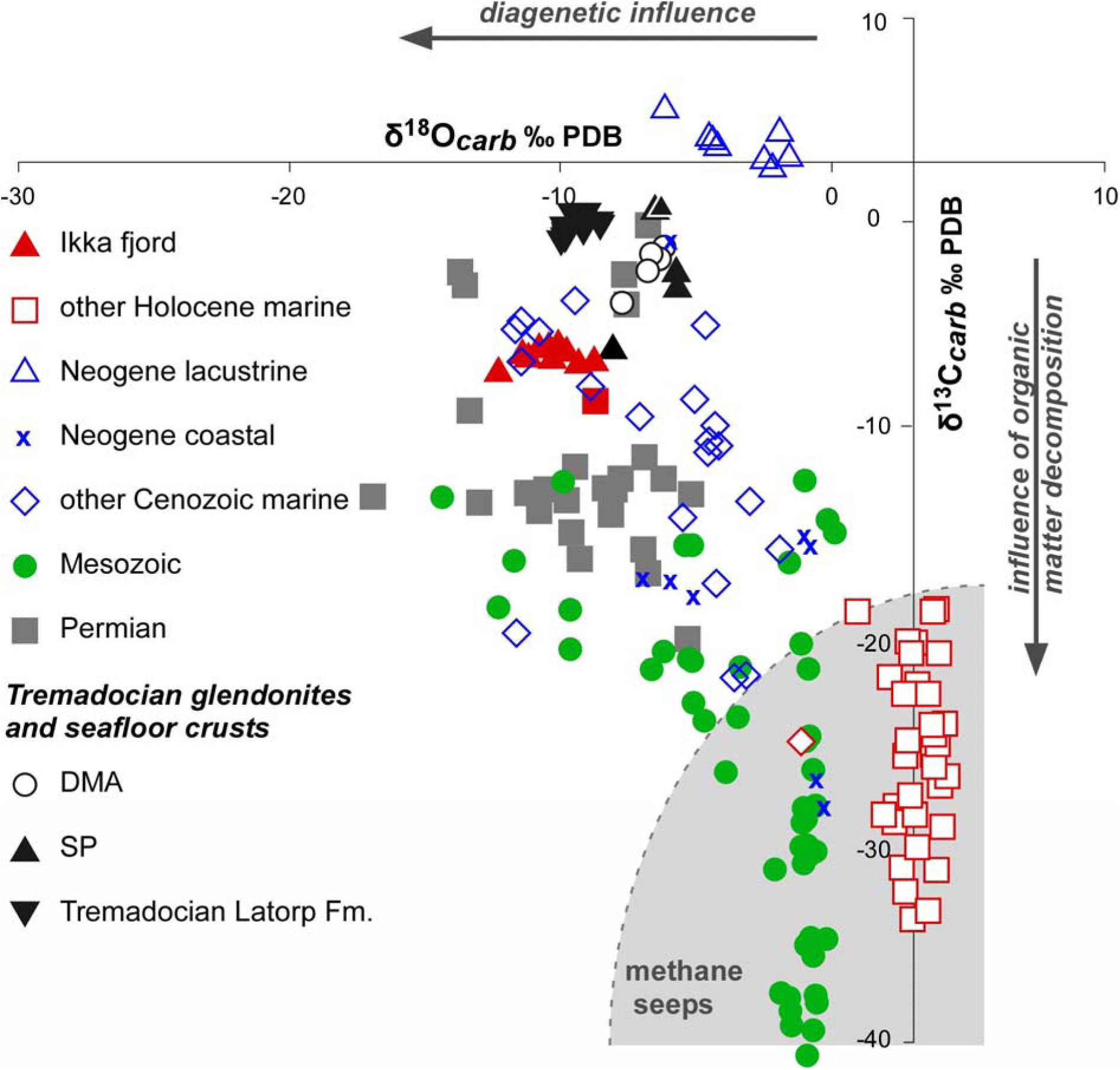
Stable isotope dataset plot of glendonite calcite and ikaite. Ordovician data in Repository Data; chemostratigraphy of the Latorp Formation after^14^; remaining data after ^3,10,13,16,33^.

Late Tremadocian to Floian glendonite-free limestone interbeds sampled in the Latorp Formation (Jämtland, Sweden), a laterally equivalent of overlying glendonite free glauconitic sands of the Leetse Formation (Supplement Figs 2–4), have yielded S^13^C values heavier than those of the study glendonites (ranging from +0.27 to −0.73‰)^14^ but lighter δ^18^O signatures (from −8.29 to −9.89 ‰)^14^, reflecting a stronger diagenetic overprint of the succession in the glendonitic-free depocentre of the Baltoscandian Basin.

Oxygen isotopes derived from glendonites should not be used for palaeotemperature purposes. As the ikaite → calcite transition results in a 68.6% volume loss, related to the release of structural water from the original ikaite crystal^12^, the resulting glendonite is controlled by pseudomorphic transformation into calcite and subsequent porosity occlusion by calcite mosaics, with different oxygen and carbon isotopic compositions than the original ikaite^9^.

## Isotope data from contemporaneous conodonts

Biogenic phosphate of three conodont samples from the glendonite-bearing Orasoja and Toolse members of the Toolse 555 drill core, in the vicinity of Kunda town, where analysed for oxygen isotopes (Fig. 1b, Supplementary Fig. 3). δ^18^O values obtained from the corresponding levels A2 (+14.4%o; *C. angulatus* Zone), B3-4 (+14.4‰, *C. angulatus* Zone) and C (+14.7‰; *P. deltifer pristinus* Zone) from this core (levels after^15^) were homogeneously low (see Repository Data).

The absolute temperatures derived from the δ^18^O values vary according to different equations: the two δ^18^O values (both 14.4‰) obtained from the *C. angulatus* Zone calibrated to NBC 120c (V-SMOV) = 21.7‰ translate 49.9 °C^16^ and 48.1 °C^17^ (V-SMOV = −1). The value based on selected *Paltodus* material from the lower *P. deltifer* Zone (*P. d. pristinus* Subzone), calculated with the same equations, translates to 48.6 °C and 46.8 °C, respectively.

## Constraining ikaite vs. δ^18^O_[phosphate]_ for the reconstruction of Tremadocian climate

Glendonites are not informative of sedimentary environments. They have been reported from lacustrine and littoral to bathyal (~ 4000 m) environments, springs, melted sea ice and even caves^13^. Generally, ikaite precipitation is favoured by elevated alkalinity, dissolved phosphate in pore waters, and its formation is often associated with organic-rich marine sediments, where methane oxidation can take place.

The occurrence of glendonite rosette clusters in calm-water clayey substrates is considered as one of the most reliable palaeotemperature indicators of near-freezing conditions^3, 18^. The preservation of lag intervals, rich in glendonitic-derived clasts, associated with symmetrical ripples marks and microbially induced sedimentary structures (Figs 2d, i), points to the episodical influence of storm-induced processes reworking these calm-water substrates that served for ikaite nucleation.

In our case study, massive precipitation of ikaite is in obvious discrepancy with the palaeotemperature interpolations suggested by the stable oxygen isotope signatures obtained from contemporaneous biogenic calcite (brachiopods) and apatite (linguliform brachiopods and conodonts) sampled in subtropical substrates suggesting higher temperatures (up to 40°C) for both the seafloor and the water column^19, 20^.

Isotope data from carbonates of subtropical Laurentia^21^ show high δ^18^O values across the Furongian–Tremadocian transition, depleted values (reflecting some strong warming) into the uppermost angulatus/lowermost *manitouensis* Zone (coeval with the basal *deltifer* Zone in Baltica), followed by a positive shift starting in the lower *manitouensis* Zone and continuing into and throughout the Floian with minor fluctuations through the late Floian^21^. This rise in δ^18^O values reflects some long-term cooling starting at a level comparable to the lowermost *deltifer* Zone in Baltoscandia^22^. Although not supported by oxygen isotopes from bioapatite (see Repository data), the scale of the relative negative-positive shift of more than 2‰ in δ^18^O during the lower Tremadocian *fluctivagus/angulatus* interval is comparable with glaciation vs. deglaciation turnovers recorded during the past 110,000 years ^23^.

Isotope data from the East Baltic require some explanation. Unusually low δ^18^O values from biogenic (conodont) phosphates of the glendonite-bearing Orasoja and Toolse members can be explained by either significant freshwater influx from surrounding land or-high surface water temperatures pointing to sharp density stratification patterns of the water column. The significant differences in δ^18^O values obtained for the water column and the water/sediment interface in fossil record are rare but not exceptional. A close analogy can be found in unusually light δ^18^O values, obtained from some shark teeth across the Palaeocene–Eocene transition in the area of North Sea^25^. In both cases the observed paradox can be explained by significant decrease in surface water salinity caused by increased fresh water fluvial discharge and sharp stratification of the water column.

A high-latitude present-day analogue is freshwater influx of Siberian rivers, bearing extremely low δ^18^O values, into the Arctic shelves^26–27^. Due to the high latitude position of Baltica in the Tremadocian, low δ^18^O values of freshwater runoff into the Baltic epeiric seas can be expected, because the oxygen isotopic yielded by rivers is drastically controlled by latitude^28–29^.

Further evidence came from the occurrence of free floating planktonic green algae *Botryococcus,* which occur in Sweden through the Alum Shale Formation, including its uppermost Tremadocian part. These algae are considered as indicators of at least freshwater influence and are common in lacustrine environment^30^.

Palaeotemperature interpolations derived from oxygen isotopes in biophosphates should take into account possible secular changes in the oxygen isotope composition of Phanerozoic seawater^31^, which could be responsible for apparently too high scores of inferred sea-water temperatures, based on different present-date-based formulas.

Finally, long-term episodes of thermally stratified water masses have been proposed for the Lower Ordovician Baltoscandian Basin to explain the widespread development of kerogenous shales rich trace-metal ore deposits and distinct separation of pelagic fossil taxa (such as graptolites and conodonts) representative of depths above and below the thermocline.

## Concluding remarks

The massive ikaite precipitation in the Tremadocian black shales of North Estonia and the adjacent St Petersburg region of Russia, which lasted ca. 5 m.y., is exceptional and the first ever recorded for the entire Cambrian-Carboniferous sedimentary record worldwide. It took place in a shallow, epeiric sea located in temperate southern latitudes, and was associated with transgressive and high patterns of organic productivity conditions. This event indicates near-freezing conditions in bottom waters over the region.

The isotopic data of the reported glendonite are not pristine, but represent a mixture of pseudomorphic transformation into calcite and occlusion of the porosity related to the ikaite → glendonite transformation by early-diagenetic calcite. As a result, they are not valid for palaeotemperature interpolations. In contrast, δ^18^O_[conodont phosphate]_ values are currently considered as pristine from widespread conodont taxa living in the water column and fossilised in glendonite-bearing strata. However, the low δ^18^O values derived from conodont apatite, which interpolate to high temperatures (>40°C), comparable to those reported in contemporaneous calcitic brachiopods from shallow-water subtropical settings, were biased by (i) the onset of thermal stratification in the semi-closed Baltoscandian Basin, probable (ii) freshwater influx and (iii) secular changes in the oxygen isotope composition of Phanerozoic seawater. The thermal/saline stratification of the Baltoscandian water column should have played an important role in moderating subpolar climates and reducing latitudinal gradients in the Tremadocian times.

## Supporting information

**Supplementary Information** is linked to the online version of the paper at www.nature.com/nature.

## Acknowledgements

Financial support for this work was provided by project CGL2017-87631-P from Spanish MINECO. OL acknowledges support by the Estonian Research Council (grant PUT 378) and Deutsche Forschungsgemeinschaft (DFG project LE 867/8-1 and 8-2), Research of MGP supported by Golestan University Research Grant 6762. Research of LEH is supported by the Zhongjian Yang Scholarship from the Department of Geology, Northwest University, Xi’an. Zhifei Zhang sincerely acknowledges the National Natural Science Foundation of China (41425008, 41621003 and 41720104002) and the 111 Centre (D17013) for the continuous financial support for the palaeobiology group in Xi’an. This paper is a contribution to IGCP 653 project.

## Author Contributions

LEP, JJA, MGP and AVD designed the study; HB, LEH, AVD, LEP, ZhifeiZ, ZhiliangZ and MGP contributed directly in specimen sampling and field documentation; JJA performed petrographical, geochemical and isotope studies of glendonite samples; PM and OH responsible for conodont biogenic phosphate sampling and its preparation for analysis; OL and MJ performed oxygen isotope analyses and discussed outcomes; LEP, JJA and LEH analyzed data and wrote the paper; MGP, HB and ZhiliangZ conducted illustration preparation. All authors discussed results and commented on the manuscript.

## Author Information

Reprints and permissions information is available at www.nature.com/reprints. The authors declare no competing financial interests. Readers are welcome to comment on the online version of this article at www.nature.com/nature. Correspondence and requests for materials should be addressed to…

## METHODS

The analysed antraconites were sampled in the Türisalu Formation (*Paltodus deltifer pristinus* Zone) of a costal section in vicinity of Udria, North Estonia (acronym **DMA**), the Koporiye Formation (*Cordylodus angulatus* Zone) close to Yurtsevo village, and the St Petersburg region (**SP**). The greatest concentration of antraconites occurs within a bed, 1 m thick, at Yurtsevo and Chernetskaya sections (Fig. 1b, Supplement Fig. 2; see also GPS settings in Repository Data).

Samples were petrographically characterized using a combination of methods, including transmitted light microscopy with thin-sections stained by Alizarin Red S and Potassium Ferricyanide, scanning electron microscopy (SEM at National Museum of Natural Sciences, Madrid) operating in back-scattered electron (BSE) image and energy dispersive X-ray (EDS) analysis, and separate cold cathodoluminescence microscopy (CL at Spanish Geological Survey, IGME-Tres Cantos, Madrid). Analytical results of back-scattered electron imaging and EDS analyses display an error of ±5 to 7%. The qualitative mineralogical composition of some complex samples was determined by the X-ray powder diffraction method. Interpretation of cementation history was made by distinguishing cement types based on colour, brightness, luminescence patterns, cement morphology and cross-cut relationships. Complex zonation of cements (revealed by CL, BSE and EDS) allowed correlation of cement zones between samples. Major, trace, and rare-earth elements were determined using X-ray fluorescence and inductively coupled plasma mass spectrometry (ICP-MS at AcmeLabs, Canada). Precision for major, trace and rare elements is usually better than 2%, 5–10% and 3–7%, respectively.

Stable isotopes of oxygen and carbon for whole-rock carbonates were removed by dental drill under a binocular microscope and analysed at Erlangen University. Carbonate powders were reacted with 100% phosphoric acid at 75°C using a Kiel III carbonate preparation line connected online to a ThermoFinnigan 252 mass spectrometer. All values are reported in permil relative to V-PDB by assigning a δ^13^C value of + 1.95‰ and a δ^18^O value of −2.20‰ to NBS19. Reproducibility was checked by replicate analyses of laboratory standards and is better than ± 0.06 ‰ (1 std.dev.).

Chemical conversion of the phosphate bound in conodont apatite into trisilverphosphate (Ag_3_PO_4_) was performed following Joachimski et al. ^34^ method and subsequent oxygen isotope analyses were performed using a TC-EA (high-temperature conversion-elemental analyzer) coupled online to a ThermoFinnigan Delta V Plus mass spectrometer. 0.2 to 0.3 mg Ag_3_PO_4_ was weighed into silver foil and transferred to the sample carousel of the TC-EA. At 1450 °C, the silver phosphate is reduced and CO forms as the analyte gas^35^. CO was transferred in a helium stream through a gas chromatograph via a Conflo III interface to the mass spectrometer and values are reported in ‰ relative to VSMOW. Samples as well as standards were measured in triplicate, measurements were calibrated by performing a two-point calibration^36^ using NBS 120c (21.7 %o) and a commercial Ag_3_PO_4_ (9.9 ‰). A laboratory standard was used as a control standard and processed together with the samples. All standards were calibrated to TU1 (21.11 ‰) and TU2 (5.45‰^35^). External reproducibility, monitored by replicate analyses of samples was ± 0.14 to ± 0.29 ‰ (1 σ).

## EXTENDED DATA

**Fig. 1.**
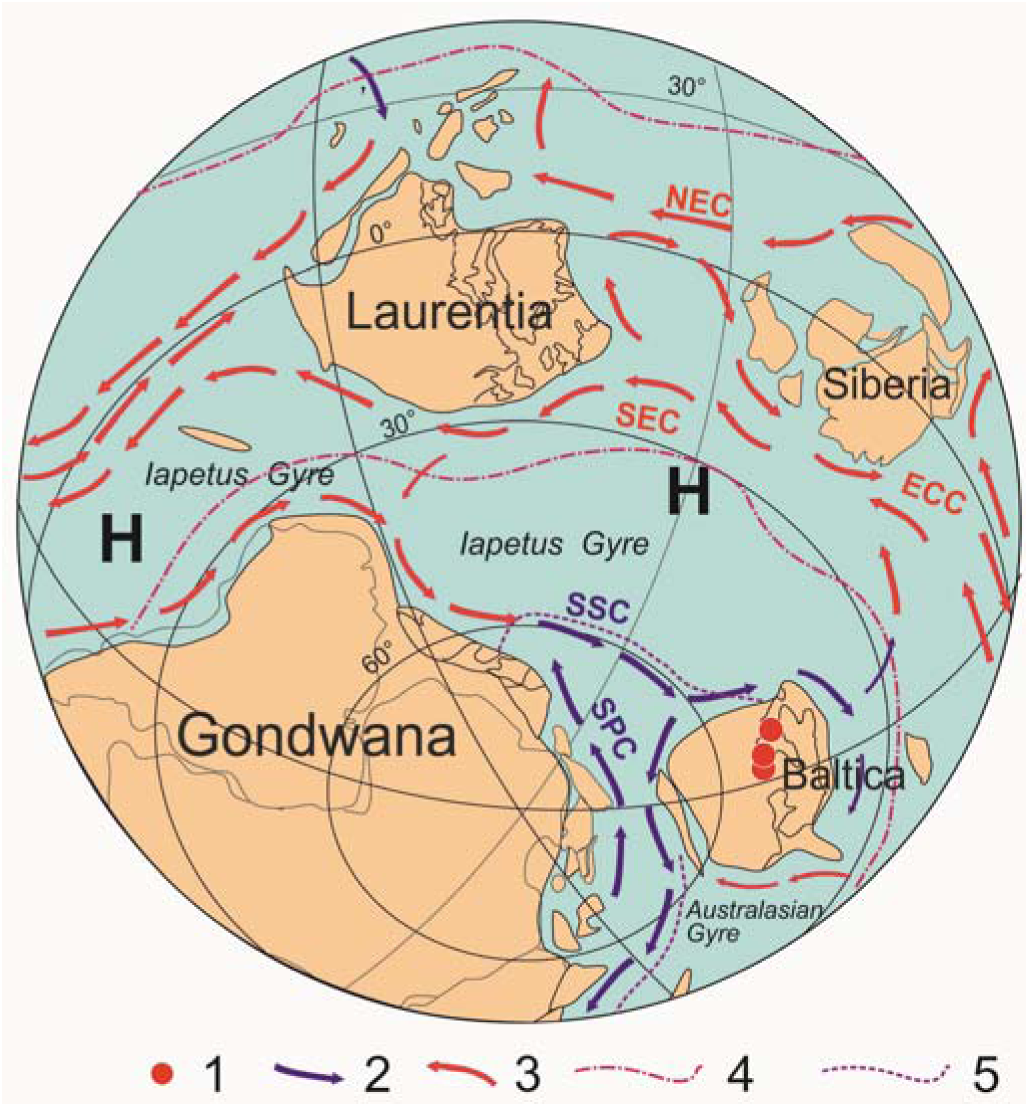
Schematic palaeogeographical reconstruction for early Tremadocian of South Hemisphere, including inferred oceanic circulation (strongly modified after Wilde^45^). Legend: 1, areas of ikaite precipitation; 2, cold water currents; 3, warm water currents; 4; Tropical Convergence; 5, Polar Convergence; NEC, North Equatorial Current; SEC, South Equatorial Current; ECC, Equatorial Counter Current; SSC - South Subpolar Current; SPC - South Polar Current; H, subtropical high pressure zones; K, Kolguyev Island; T, Timan – Pechory Region; CA, Central Africa.

**Fig. 2.**
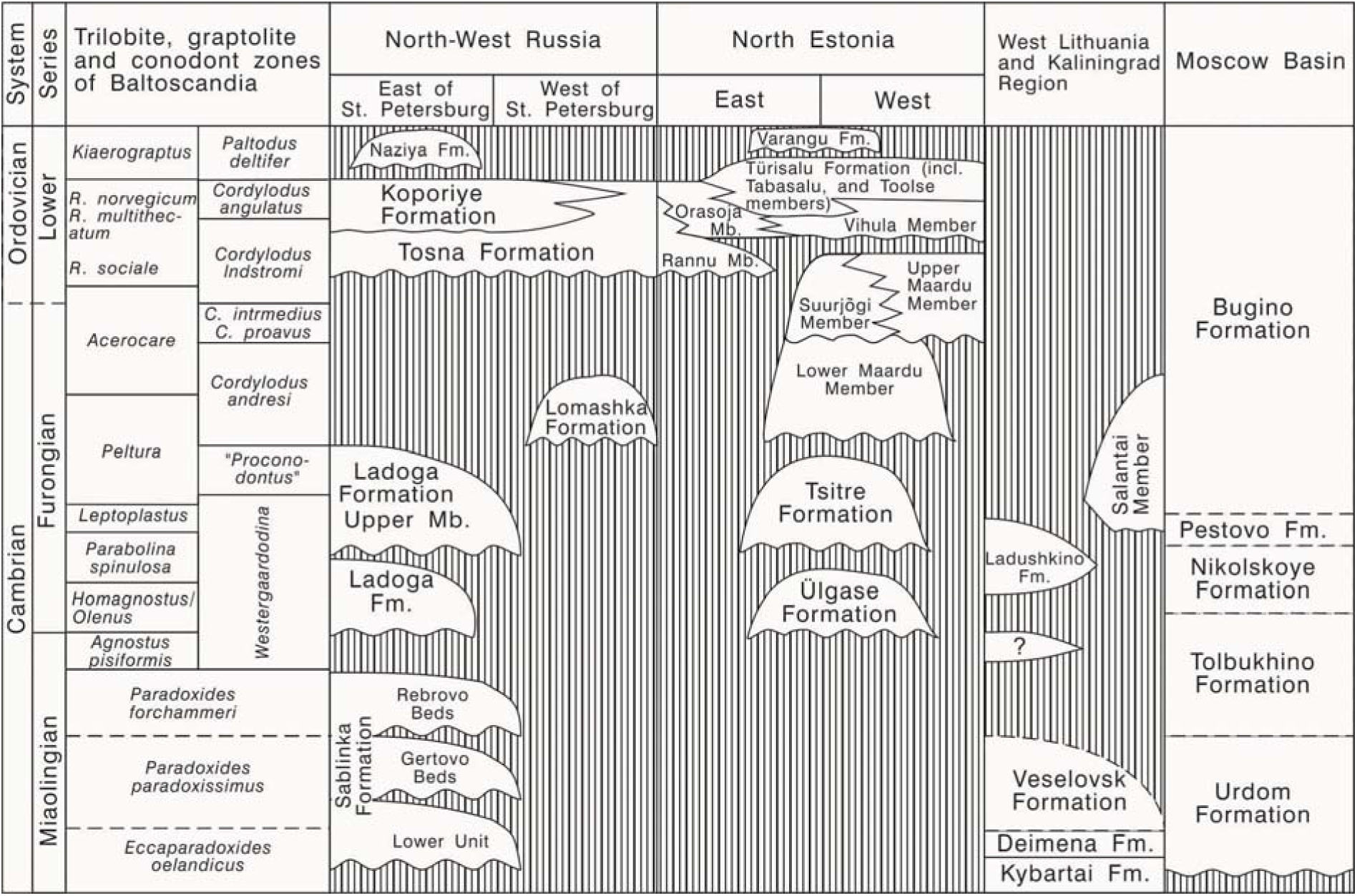
Revised lithostratigraphical correlation chart for the Cambrian (Miaolingian) - Ordovician (Tremadocian) of North Estonia and European Russia.

**Fig. 3.**
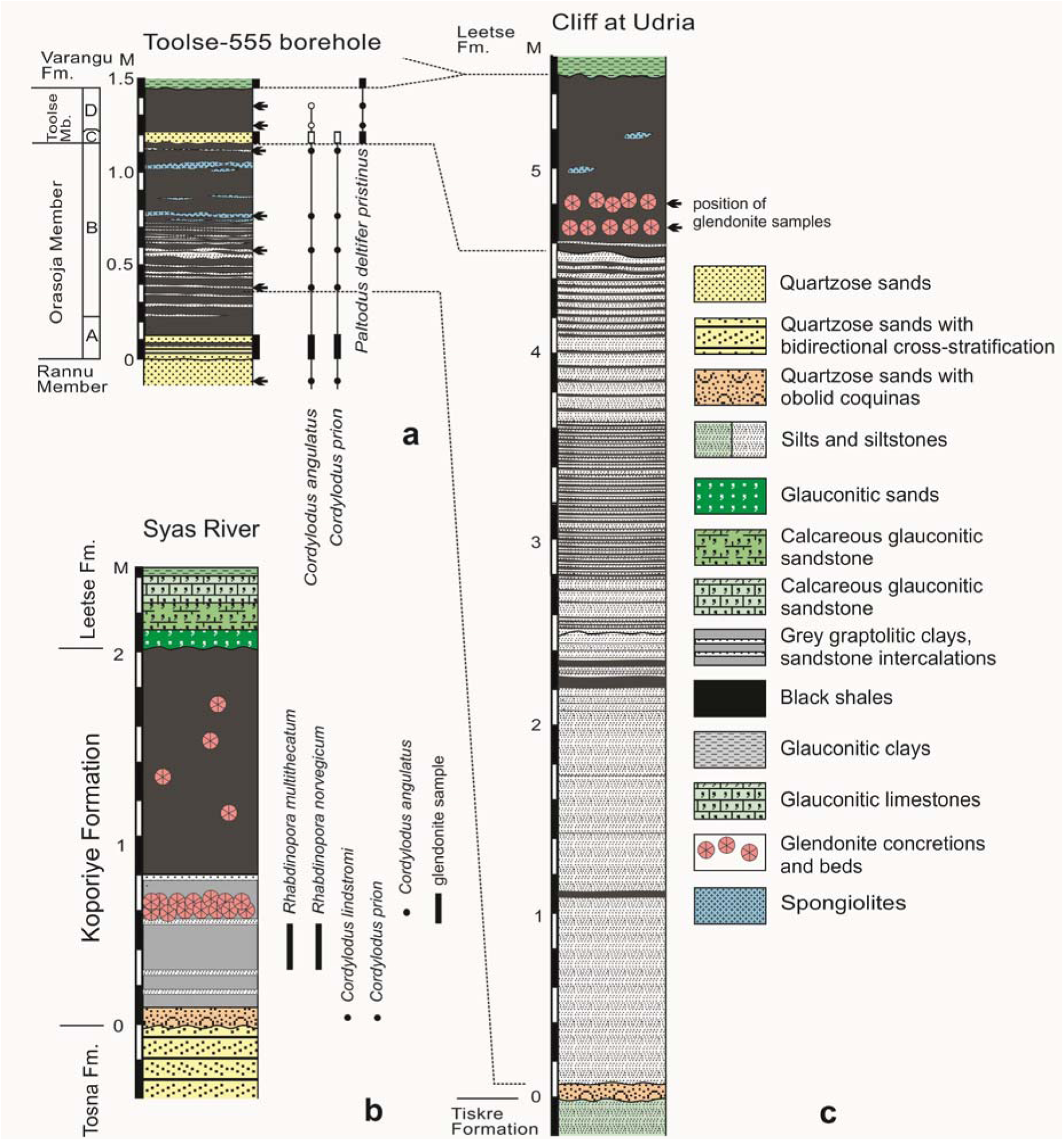
Lower Tremadocian stratigraphical logs sampled for glendonites and conodont biogenic phosphates; **(a)** drill core of T-555 borehole in vicinity of Kunda (Fig. 1b) ^15^, A, B, C show stratigraphical intervals of conodont samples used in the oxygen isotope analysis; **(b)** Koporiye Formation in vicinity of Kolchanovo and Yurtzevo, showing glendonite occurrences; **(c)** Orasoja Member of Kallavere Formation and Toolse Member of Türisalu Formation showing setting of glendonite samples.

**Fig. 4.**
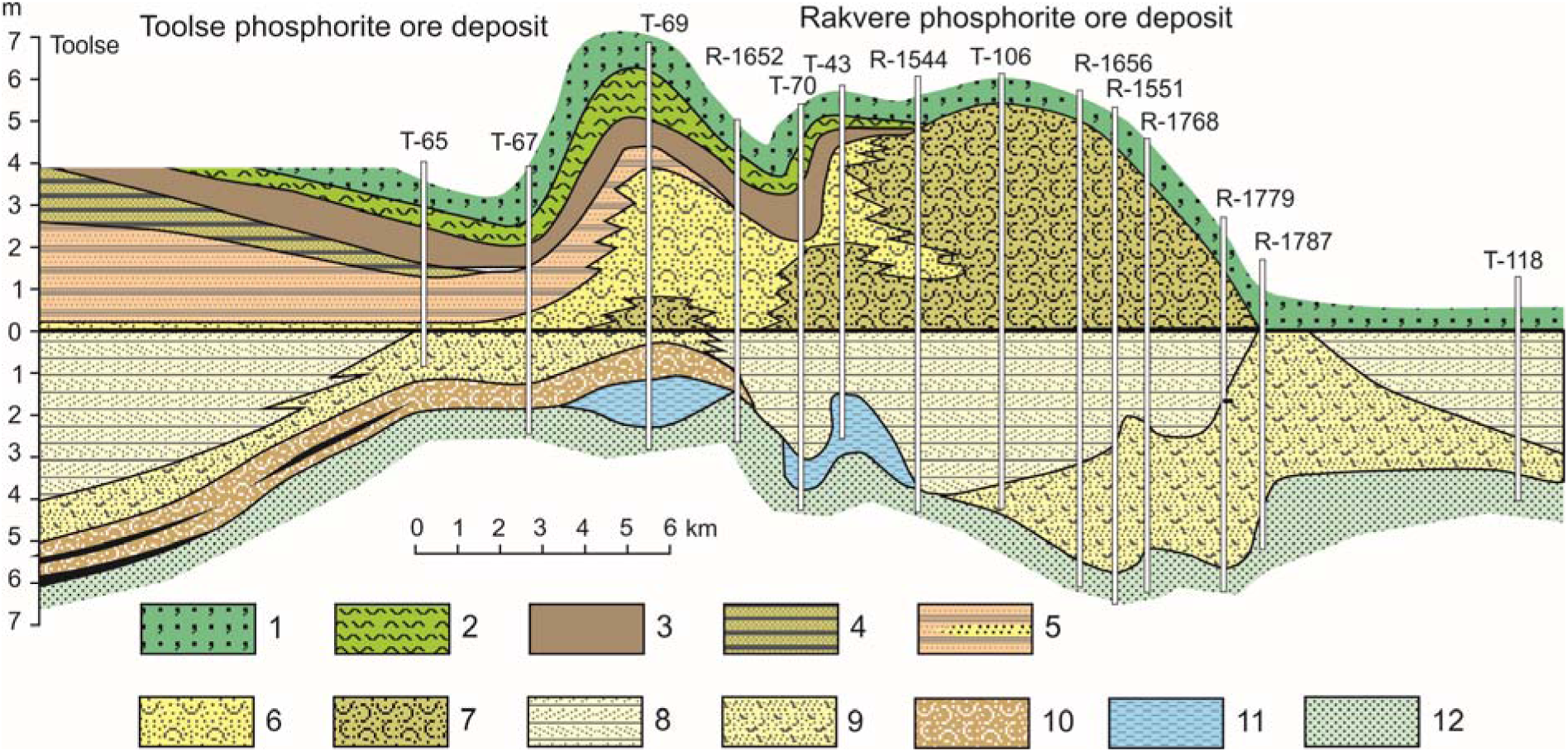
Geological cross section through Furongian-Tremadocian deposits in North Estonia. Legend: 1, glauconitic sands of Leetse Formation (Floian); 2, glauconitic clays of Varangu Formation (Tremadocian); 3, black shales of Türisalu Formation (Tremadocian); 4–12, Kallavere Formation (upper Furongian-Tremadocian); 4, Orasoja Member of black shale interbeds (P_2_O_5_<3%); 5, Vikhula Member of fine grained sands with reworked obolid shells and subsidiary black shales (P_2_O_5_=1-3%); 6, 7, Rannu Member of quartzose sands (P_2_O_5_ =3-9%) and detrital phosphoritic sands (P_2_O_5_>9%); 8, Suurjõgi Member of quartzose sands often with bidirectional cross-lamination (P_2_O_5_=3-9%); 9, 10, Maardu Member of silty sands with subsidiary black shales (P_2_O_5_=1-3%) and coquina accumulations (P_2_O_5_>18%); 11, Ülgase Formation (lower Furongian) of silts and silty clay; 12, Tiskre Formation (Cambrian Series 2). Zero datum level coincides with the inferred position of the Cambrian–Ordovician boundary^5^ (modified).

**Figure 5.**
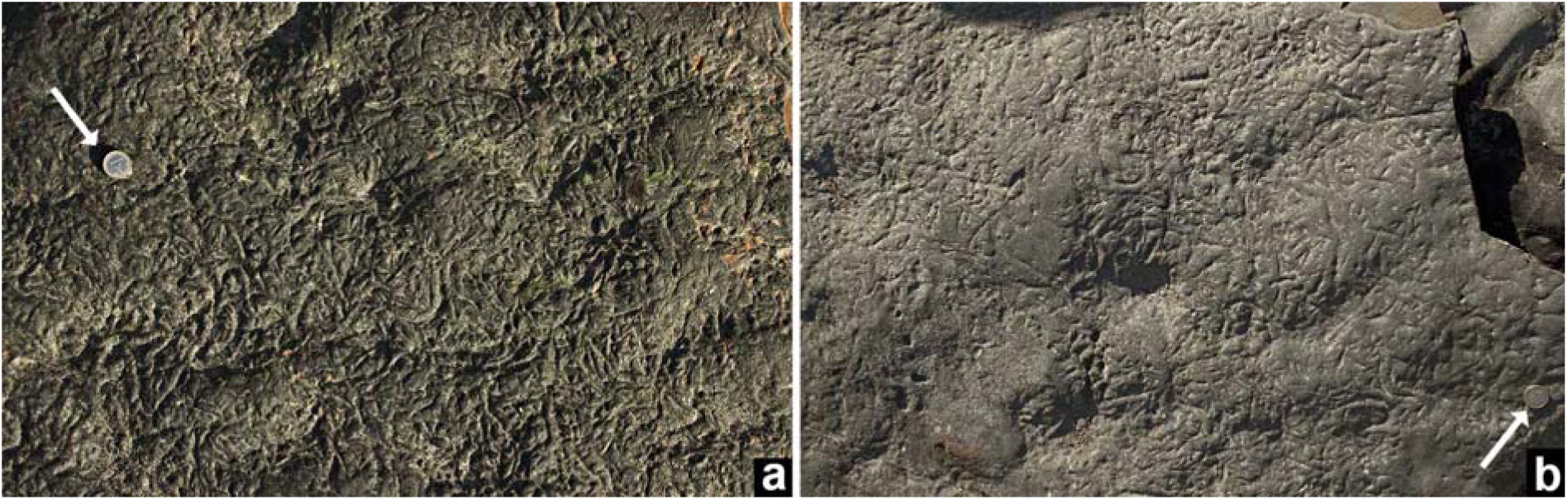
View of upper-bedding surfaces resembling *Kinneya* wrinkle structures showing meandering and partly interfering, flat-topped to rounded crests and intervening grooves and pits; **(a)** with higher and **(b)** lower density patterns; base of Türisalu Formation at Pakri cape, Estonia; scale arrowed.

**Figure 6.**
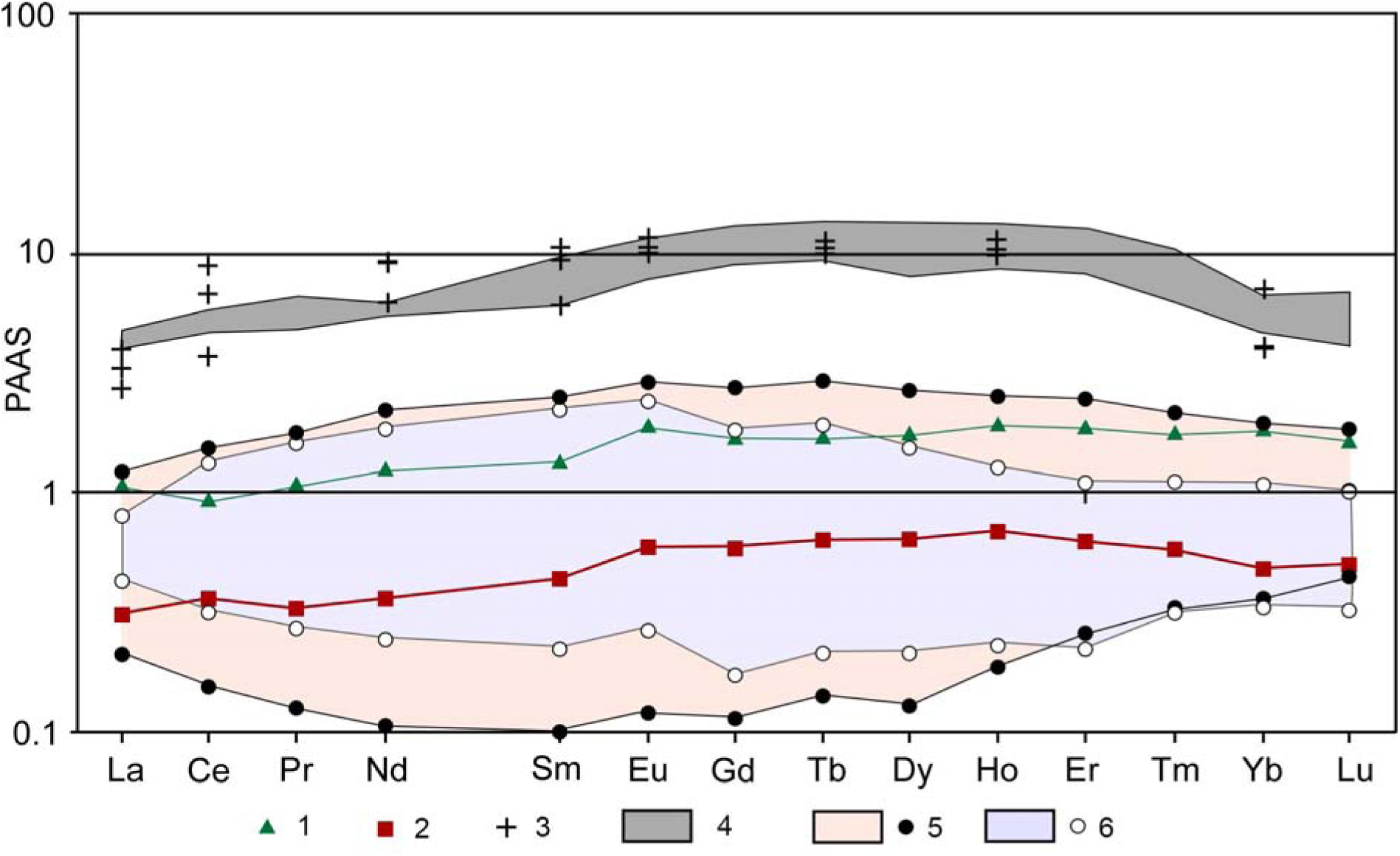
Shale-normalized REE distribution patterns for Tremadocian black shales and glendonites from East Baltica. Legend: 1, black shales of Koporiye Formation; 2, glendonite average SP1-4; 3, bioapatite (obolids) from sands of Ladoga Formation and Maardu Member^83^; 4, bioapatite (obolids) from sands of Kallavere Formation^95^; 5, black shales of Tabasalu Member from Pakri^75^; 6, Black shales of Toolse Member from Saka^75^ (PAAS shale values^87^).

Table of chemical analyses.

| | East Baltic Stage | Conodont Zone | $\delta^{18}\text{O}$<br>(‰VSMOW) | 1 std. dev. |
| --- | --- | --- | --- | --- |
| OL-A2 | Pakerort Stage | *Cordylodus angulatus* | 14,42 | 0,14 |
| OL-B 3-4 | Pakerort Stage | *Cordylodus angulatus* | 14,44 | 0,26 |
| OL-C | Pakerort Stage | *Paltodus deltifer*<br>(*P. d. pristinus*<br>Subzone) | 14,66 | 0,29 |

