## Supplementary material for "Glendonite occurrences in the Tremadocian of Baltica: first Early Palaeozoic evidence of massive ikaite precipitation at temperate latitude"

### SUPPLEMENTARY INFORMATION

#### Geological setting and stratigraphy

The Tremadocian Age was at the dawn of a spectacular increase in marine biodiversity at all sub-phylum taxonomic levels, known as the Great Ordovician Biodiversification Event. Different sedimentological and geochemical indicators apparently point to global greenhouse conditions, accompanied by bottom oxygen-depletion in epeiric seas. Atmospheric $CO_2$ levels were likely 10 to 16 times preindustrial levels[37], the South Pole was located at NW Africa, probably, under a shallow sea[38], solar luminosity was 3.5–5.0% less than modern values[39-40], and terrestrial plants and consequent climate feedbacks were effectively absent[41-43]. Climate simulations point to equatorial temperatures about 30–35ºC during ice-free portions of the Ordovician[44-45].

At that time Gondwana supercontinent occupied a vast space from the South Pole to equatorial latitudes, although significant area within the polar circle was exposed above sea level. Baltica continent located in a close proximity of the North African sector of Gondwana with the Uralian margin facing south, completely within the temperate latitudes and was exposed to the cold oceanic currents from its southern and western, Caledonian side (Supplementary Fig. 1). The influx of cold, saline oceanic waters from the Caledonian Baltic margin is supported by the signs of strong westerly directed (modern coordinates) bottom currents documented across the area[46]. Two other major continents, Laurentia and Siberia located on both sides of the equator engulfed by the global system of warm equatorial currents; thus they were isolated from significant impact of secular climatic changes in the South Hemisphere. This arrangement of major land masses suggest existence of two major wind-driven ocean gyres in southern temperate latitudes[47].

Oxygen isotopic values derived from Early Ordovician calcitic brachiopods (exclusively from low latitudes) are consistently low (–8.2 to –11.1 ‰VPDB)[48-49] suggesting a combination of warm temperatures, a pervasive diagenetic overprint, and very low Early–Ordovician seawater $\delta^{18}O$ values[48-51]. Tropical seawater palaeotemperatures calculated in West Laurentia[20] from micrite $\delta^{18}O$ values and assuming ice-free seawater, suggest tropical seawater temperatures averaged 42ºC, whereas $\delta^{18}O$ values of co-occurring biogenic apatite yield estimates averaging 37ºC. The phosphatic values are interpreted as less diagenetically affected than carbonate values[20] and the presence of linguliform brachiopods and conodonts in mid- to high-latitude platforms have recently multiplied the application of $\delta^{18}O_{\text{[biogenic phosphate]}}$ values as reliable palaeotemperature indicators [19, 52-61]. However, the different provenance of the analysed biogenic material (benthic brachiopods vs. pelagic conodonts) is not currently taken into account, and Tremadocian seafloors are currently interpolated as representative of warm surficial conditions worldwide. Furthermore, recent developments in biomineralisation studies clearly demonstrate that biogenic hydroxyapatite is metastable outside the animal body[62-63], which creates new challenges in application of biogenic phosphates for interpretation of isotope analysis outcomes and requires better understanding of their chemical taphonomy.

This paper aims to provide a new high-temperate-latitude palaeoceanographic scenario based on the coexistence, in the Tremadocian of East Baltica, of (i) climate-sensitive indicators of bottom waters close to the freezing point (glendonites) and (ii) geochemical indicators of warmer temperatures for the water column.

In Eastern Baltica, the Furongian–Tremadocian sedimentary succession (Tsitre and Kallavere formations in North Estonia; Ladoga, Lomashka and Tosna formations in the St Petersburg region of Russia; Supplementary Figs 2-4) comprises condensed, unlithified, cross-laminated quartzose sands, silts and clays with subsidiary black shale interlayers that increase in thickness upsection. The total thickness of this interval does not exceed 10 m, except in western Estonia, where it approaches 15 m. Despite the superposition of condensed phosphate and ironstone crusts and considerable hiatuses, the East Baltic biostratigraphic succession is remarkably complete through the Cambrian–Ordovician boundary interval: all Baltic conodont zones within the interval from *Cordylodus andresi* to *Paroistodus proteus* are documented[4]. Therefore, a continuous black shale deposition occurred within the interval of *Cordylodus angulatus* to *Paltodus deltifer pristinus* zones, marking the peak of the Early Tremadocian marine transgression. The basin was rimmed to the south (recent coordinates) by a chain of low islands and associated shoal complexes (e.g. Rannu Member of the Kallavere Formation; Fig. 1) traceable over 600 km from Pärnu at the west to Volkhov at the east (Fig. 1). These shoals and bars contain high concentrations of allochthonous obolid coquinas, re-deposited from Furongian bars[5] and represent an economically significant source of phosphorous (only the Rakvere Phosphorite Ore Field contains roughly 700 million tons of biogenic $P_2O_5$[64]). Some remnants of Furongian bars (e.g. Maardu and Toolse phosphorite ore deposits; Fig. 3) were preserved from complete erosion due to fast sea-level rise, which placed them below the wave influence. The black shales appear as layers up to 10 cm thick intercalating with fine-grained, phosphoritic sands in the lower part of the Orasoja Member of the Kallavere Formation, which is considered as part of a shoal complex[65]. Black shale beds in the upper part of the unit in the Orasoja section are rife of glendonites[65].

A continuous deposition of black shales in the Baltoscandian Basin commenced at the beginning of the *Cordylodus angulatus* Zone and represents the transgressive phase of Black Mountain Eustatic Event[66] and the Stonehenge Transgression of Baltica[22]. An estimated magnitude of the sea-level rise was about 40–60 m, bringing the seafloor close to the storm wave base level[67]. However, shallowing-upward trends episodically allowed organic-rich mud deposition within the photic zone and under the storm and wave action[68].

Throughout Cambrian times, Baltica lay at southerly latitudes (~ 30–60°S[32]) and was geographically inverted: present-day southern Baltica (including Scania) faced the equator and NW Baltica West Gondwana. A key palaeomagnetic datum from a limestone in the Furongian Alum Shales, sampled in the Forsemölla – Andrarum area, places Scania at ca. 35°S[69] (for other views see[70-71]).

**Sedimentary and early diagenetic environments**

The Tremadocian Türisalu and Koporye formations accumulated in a shallow-water inland extension of an exceptionally flat-floored epeiric sea, the Baltoscandian Basin. As stated above, the basin was episodically rimmed to the south (recent coordinates) by a chain of low islands and associated shelly shoal complexes, while its depocenter since Furongian shows continuous deposition of black shales, which spread towards the shallows in the early Tremadocian. Here black shales are ranging from the uppermost *Cordylodus lindstromi* to *Paltodus deltifer pristinus* conodont zones[4]. Their top is disconformably overlain by the organic-poor grey clays and glauconitic sands of the Varangu Formation.

The discontinuous chain of low islands that fringed the Baltoscandian Basin was rimmed by Furongian biogenic phosphate-rich shoal systems, which were flooded and reworked during the Tremadocian marine transgression (Supplementary Fig. 4). The shore line moved several tens kilometres south[5]. It was rimmed by shoal complexes enriched in allochthonous biogenic phosphate (e.g. Rakvere phosphorite ore deposit). That old land was flooded completely only in the late Floian time.

These phosphoritic (weight percent $P_2O_5$>18%) bars played a significant source for the increasing phosphate pollution of the oritic crusts along the margiTremadocian water column, especially when high alkaline dysaerobic conditions developed at the sediment-water interface[5, 68]. The latter conditions are supported by significant deposition of thin phosphns of the black shale depocentre (Fig. 1a; Supplementary Fig. 4)[72]. These phosphorites contain *Rhabdinopora* graptolites[5], which leave no doubts that they were synchronously deposited with the black shales of the Türisalu Formation.

The kerogenous and metalliferous black shales of the Türisalu and Koporiye formations display shallowing-upward parasequences, up to 7 m thick. Massive to laminated black shales are topped by siltstone interbeds rich in cross- and wavy-lamination, symmetric ripple marks, centimetre-scale lag deposits with grading, scouring surfaces and episodic record of burrowing and spiculites [65, 67-68]. The parasequences are associated with vertical decimetre-scale redox-sensitive trace metal shifts[73-75] reflecting metal sequestration related to contemporaneous fluctuations in sedimentation rate. In NE Estonia, the Toolse Member (Türisalu Formation) becomes thinner and consists of a centimetre-thick alternation of siltstones and black shales containing authigenic carbonate and sulphide mineralisations[76-77]. In Scania, the influence of relative high-order sea-level fluctuations is recognized by the vertical stacking pattern of different claystone-dominant facies associations of the Alum Shale[78]. In the parasequences of the Türisalu and Koporiye formations, glendonites are relatively common in the lower massive-to-laminated black shales and the upper black shale interbeds of the upper part. The latter contain scattered centimetre-thick lag deposits rich in clasts derived from glendonites and seafloor crusts.

Oxygen content was also variable, ranging from temporary oxygenated conditions above the sediment-water interface (supported by the episodic occurrence of ichnofossils, often preserved as a result of pyrite infill, and thin, lens-like layers of spiculites and associated acrotretid brachiopods[5, 68]) to dysaerobic, high alkaline conditions proved by the enrichment in redox-sensitive metals[8], including molybdenum, uranium and vanadium. Oxygenation episodes are recognized by the sudden development of metazoan colonization, burrowing and development of microbially induced sedimentary structures[68, 79]. Microbial mats were then able to stabilise the seafloor, protecting against erosion and increasing the cohesiveness of the marine substrate. It is noticeable the local abundance of *Kinneya*-type wrinkle structures, considered as subsurface structures developed on a clayey substrate underneath biofilms and mats[80-81].

The Furongian–Tremadocian Nd isotopic signatures show median values of the $\varepsilon$Nd(t) within the range from –7.0 to –8.0 for the whole Baltoscandian Basin[82]. These data were probably close to the original signatures of the adjacent Iapetus oceanic water masses, suggesting free exchange: there are no signs of the high negative $\varepsilon$Nd(t) valued characteristic of old cratons. Therefore, a significant part of the Baltic continent, including the Fennoscandian Shield, was covered by epeiric seas. Sedimentation rates of the East Baltic black shales were extremely low, below 10 mm per millennium and approaching those of pelagic clays in present-day oceans[70].

**REE signatures**

A geochemical comparison in rare earth elements (REE) is made between Furongian and Tremadocian brachiopod shells and conodonts[83], the Tremadocian black shales of Estonia[77], completed with new analyses from the Koporiye Formation of the St Petersburg area, and the above-reported glendonites and seafloor crusts (see Repository data), as a geochemical clue to constrain recycling of Furongian apatite in the Tremadocian water column.

X-ray fluorescence (XRF) analysis shows that the analyzed Furongian biogenic phosphorites consist of francolite (carbonate fluorapatite) with and $P_2O_5$ content from 9.07 to 35.75 wt% and CaO contents ranging from 13.22 to51.51 wt%, whereas detrital debris ranges from 0.74 to 62 wt% $SiO_2$ and 0.19 to 2.32 wt% $Al_2O_3$. This contrasts with the $SiO_2$ and $Al_2O_3$ contents yielded by the Tremadocian black shale host, ranging from 35.88 to 55.11 wt% and 10 to 14.40 wt%, respectively. The total organic content (TOC) can reach up to 10.81% in the Türisalu black shales of Estonia[84], and up to 14wt% in the Tremadocian Alum black shales of Scania[85].

Inductively coupled plasma mass spectrometry analyses reveal that the total REE content is highly variable (Supplementary Fig. 6) with about $10^3$ times for isolated linguloid brachiopods (enrichment due to post-mortem adsorption of REE at the sediment-seawater interface[86]), up to >10-fold enrichment relative to average shale (represented by the PAAS shale standard[87] for shelly phosphorites, and from 0.1 to >1.0-times for black shale and glendonites. High REE concentrations in phosphates are related to their incorporation following precipitation in the marine environment and during subsequent early diagenesis[89]. A conspicuous middle REE-enriched (defined as Sm-Ho) pattern, known as the “hat pattern” and common in biogenic apatites[90-91], is recognized in the Furongian obolid accumulations. Its record is consistent with an early diagenetic origin[92] that may be associated with the reductive dissolution of iron oxides in anoxic pore waters[93]. As a result of such diagenetic scavenging, phosphorites commonly fail to retain the defining characteristics of modern (oxic) seawater REE distribution patterns, typified by progressive enrichment in the heavier REE, a pronounced negative Ce anomaly, and positive La and Y anomalies[94].

As they represent bulk analyses, REE distribution patterns do not solely represent original values but also the authigenic overprint. By comparison with published data from Tremadocian (Türisalu) black shales from Estonia[73] completed with new data from the Koporiye Formation, similar REE distribution patterns (with faint Sm-Ho enrichment) are exhibited by the black shales from the Pakri and Saka (pars) sections, but not by part of the latter section, which show a symmetric pattern with middle REE (Sm-Ho) depletion, opposite to that recorded in the phosphates. Ce and Sm-Ho depletion may be related to scavenged REE under anoxic conditions during early diagenesis. Both REE distribution patterns are mimicked by the analysed glendonite, which exhibit both middle-REE enrichment and depletion, directly controlled by early diagenetic conditions.
